# What matters to patients?: a timely question for Value Based Care

**DOI:** 10.1101/2020.01.03.893826

**Authors:** Meron Hirpa, Tinsay Woreta, Hilena Addis, Sosena Kebede

## Abstract

**Background:** Our healthcare system is moving towards patient-centered and value-based care models that prioritize health outcomes that matter to patients. However, little is known about what aspects of care patients would prioritize when presented with choices of desirable attributes and whether these patient priorities differ based on certain demographics.

**Objective:** To assess patients’ priorities for a range of attributes in ambulatory care consultations across five key health service delivery domains and determine potential associations between patient priorities and certain demographic profiles.

**Methods:** Using a *What Matters to You* survey patients ranked in order of importance various choices related to five health service domains (patient-physician relationship, personal responsibility, tests/procedures, medications and cost). Subjects were selected from two Johns Hopkins affiliated primary care clinics and a third gastroenterology subspecialty clinic over a period of 11 months. We calculated the percentage of respondents who selected each quality as their top 1-3 choice. Univariate and multivariate analyses determined demographic characteristics associated with patient priorities.

**Results:** Humanistic qualities of physicians, leading a healthy lifestyle, shared decision making (SDM) for medications and tests/procedures and knowledge about insurance coverage were the most frequently ranked choices. Privately insured and more educated patients were less likely to rank humanistic qualities highly. Those with younger age, higher educational attainment and private insurance had higher odds of ranking healthy lifestyle as a top choice. Those with more education had higher odds of ranking SDM as a top choice.

**Conclusions:** Identifying what matters most to patients is useful as we move towards patient-centered and value based care models. Our findings suggest that patients have priorities on qualities they value across key health service domains. Multiple factors including patient demographics can be predictors of these priorities. Elucidating these preferences is a challenging but a valuable step in the right direction.

## Introduction

Health systems are moving towards a Value Based Care (VBC) model of service delivery which focuses on health outcomes and cost containment.[1–6] One commonly accepted definition of value-based healthcare is “the creation and operation of a health system that explicitly prioritizes health outcomes which matter to patients relative to the cost of achieving this outcome.”[5] A related care philosophy, Patient Centered Care (PCC) also highlights the importance of addressing what matters to patients during their healthcare experience.[7–11] One challenge in the measurement of patient experience is the difficulty of differentiating among multiple overlapping terms like satisfaction, engagement, perceptions, priorities, values and preferences.[12–14] Patient preference and value can also be highly dynamic and dependent on several factors including patients’ health status, and personal characteristics such as education level.[16–19]

Despite the limitations of patient-reported measures, patient surveys can provide helpful data to identify patient preferences and values. That in turn can improve the delivery of patient-centered health services, quality of care and outcomes.[12,16,20]

Patients value both the *technical* (quality of clinical care: such as provider knowledge and skill) and the *interpersonal* (quality of communication: such as Shared Decision Making) qualities of care.[22–24] Multiple studies identify patient-provider communication to be the most important aspect of care that patients value for high-quality health care regardless of variations in socio-demographic or health characteristics of patients.[4,17,25–27] Some evidence exists that when choosing a primary care physician, the majority of patients have a strong preference for physicians of high technical quality if forced to make a tradeoff between interpersonal and technical skills.[19,28–30]

How value of the various attributes of healthcare may vary by certain patient demographics and reasons for presentation in the ambulatory primary care setting has been postulated before.[15,31] There is data that suggests low-income patients, those with a high prevalence of psychosocial problems and those feeling unwell have a preference for good communication and personal interaction when compared to their counterparts.[11,15,32] Some studies have shown that older patients are less likely to prioritize good communication[11,19] whereas other studies show that older patients at the end of life valued effective communication and trust of the provider.[32]

There is limited research examining how patients would prioritize a list of desirable attributes about specific aspects of their care, if forced to make choices. To our knowledge no study has examined patients’ priorities across key healthcare domains that we tested with concurrent assessment of demographic associations.

In this observational study, we assessed patients’ priorities for a range of attributes in ambulatory care consultations across 5 domains: patient-physician relationship, personal responsibility, tests/procedures, medications and healthcare cost and then examined potential association between patient priorities with certain demographic profiles.

## Methods

### Survey Development

We developed a 5-question survey instrument, *What Matters to You*, using 5 key health service domains: patient-physician relationship, personal responsibility, tests/procedures, medications and healthcare cost. These health service domains were previously used to determine level of shared understanding between patients and their physicians.[23,33–36] We asked patients to rank in order of importance various choices related to the 5 domains (see table 2 and Appendix). To determine whether priorities varied among subgroups of patients, we collected demographic data including age, sex, ethnicity, highest level of education and type of medical insurance.

### Participants

The subjects of this study were patients being evaluated at two Johns Hopkins affiliated primary care clinics and a third gastroenterology subspecialty clinic: Johns Hopkins Community Physicians-Remington (a primary care clinic in a suburban community in Baltimore), Johns Hopkins Community Physicians-East Baltimore Medical Center (a primary care clinic in an urban underserved community in East Baltimore) and Johns Hopkins Gastroenterology and Hepatology Clinic (a gastroenterology outpatient clinic in a suburban community 30 minutes south of Baltimore). The Johns Hopkins Institutional Review Board and the Johns Hopkins Community Physicians Research and Projects Committee approved this study. All participants were advised, verbally and in a written form, that their completion of the survey will serve as their consent to be in the research study.

### Study Design, Sample Size and Data Collection

From 7/1/2018-6/30/2019, a total of 338 patients were surveyed prior to seeing their physician in clinic. A predominant number of patients surveyed (n=298) were primary care patients. After patients were roomed for their visit, before seeing their physicians, patients were asked if they would participate in a 5-10-minutes self-administered survey designed to assess their preferences surrounding the healthcare service they receive.

### Measures

Our main outcome measures were based on the participants’ ranking of three to four important qualities under each of the five domains in the order of their personal priority.

There are 3 specific outcome measures we looked at:

1. What specific qualities under each healthcare domain were most frequently ranked as the number one choice.
2. What patients ranked as their second and third choices, as we recognized that we were ‘forcing’ patients to choose from a list of desirable attributes and wanted to assess whether there would be clear “winners”
3. Patients’ demographics as potential predictors of the most frequent top choice under each of the five healthcare domains.

### Statistical Analysis

Incomplete and erroneously completed questionnaires (n=112) were excluded from analysis. Data from the accurately completed questionnaires (n=226, 196 of which were primary care patients) were aggregated and analyzed using Excel and Stata 15.1. Some patients inadvertently received a version of the survey that had 4 choices for question four instead of 5 and 4 choices for question 5 instead of 3. Therefore, of the accurately completed surveys (n=226), an additional 53 and 95 surveys were excluded from analysis for questions 4 and 5, respectively. As a result, when calculating percent respondents for questions 4 and 5, 173 surveys for question 4 and 132 surveys for question five were analyzed.

To assess which qualities were most important for patients under each of the five domains, the percentage of respondents who selected each quality as their number one choice were determined. Since patients were forced to prioritize among a list of desirable attributes, the qualities that were ranked as the most frequent second and third choice were also determined. For questions that had greater than or equal to four choices (Questions 1 and 4), the most frequent first, second, and third choices were calculated while only the most frequent first and second choices were calculated for questions that had three choices (Questions 2, 3 and 5).

Univariate and multivariate logistic regression analyses were used to determine whether patient characteristics such as age, sex, race, education and insurance type were significant predictors of the qualities most frequently ranked as number one for each of the five domains.

During analysis, for Question 1, the choices “Kindness” and “Efforts to connect with me as a human being and not just as a patient” were combined under the heading “humanistic qualities”. For Question 2, survey option “Learn as much as I can about my condition and be actively involved in decision making” was categorized as “shared decision making”. For Question 4, survey options “I want to know exactly what I am taking and why” and “I want to understand the side effects of each medication thoroughly before accepting the prescription” were combined under the heading “shared decision making”.

## Results

Table 1 shows the distribution of participants according to age, gender, race, education level and health insurance. The mean age was 42.6 years. The study population was predominantly female (77.9%); 54% had college or post graduate degrees, 45.9% had some college or below education; and 74.1% were privately insured. There were about an equal percentage of Blacks (41.6%) and Whites (44.7%).

**Table 1.** Demographics of Overall Study Population (N = 226)

| Characteristic | N (%) |
| --- | --- |
| Age (years) | 42.6* (19-83) |
| Sex |  |
| Female | 176 (77.9) |
| Male | 50 (22.1) |
| Race |  |
| Black | 94 (41.6) |
| White | 101 (44.7) |
| Other | 31 (13.7) |
| Education |  |
| High school or less | 41 (18.1) |
| Some college | 63 (27.9) |
| College graduate | 32 (14.2) |
| Postgraduate degree | 90 (39.8) |
| Insurance |  |
| Medicaid | 24 (10.9) |
| Medicare | 27 (12.3) |
| Private | 163 (74.1) |
| Other | 6 (2.7) |

\*Mean (range)

Table 2 shows the percentage of patient respondents who ranked each quality as number one under each of the five domains. For question one assessing patient-physician relationship, “humanistic qualities^1^” (33%) was the most frequent number one choice while knowledge of the physician and ability to explain things fully were tied at 23% as the second most frequent top choice. For question number two assessing patient personal responsibility, leading a healthy lifestyle (47%) was the most frequent top choice while shared decision making^2^ (35%) and following medical recommendations (18%) were the second and third top choices, respectively. For question number three on tests and procedures, the most frequent top choice was shared decision making (50%) while wanting all tests that could be helpful (43%) and only wanting the absolute critical tests (7%) were the second and third top choices, respectively. On question four assessing medications, shared decision making^3^ (80%) was the most frequent top choice while wanting the absolute minimum medications (9%) and wanting any medication that could help (9%) were the most frequent second choice. Wanting the freedom to try alternative medicine and herbal supplements (2%) was the least frequent choice. On question five assessing healthcare cost, knowing what insurance covers (57%) was the most frequent choice while knowing what charges are for (32%) and minimizing healthcare expenditure (11%) were the second and third choices, respectively.

**Table 2.** The percentage of respondents who selected each quality as top choice.

| Patient-Physician Relationship N=226 | Percent Respondents |
| --- | --- |
| Humanistic qualities<sup>5</sup> | 33% |
| Fund of knowledge | 23% |
| Explaining things fully and in the way I understand | 23% |
| Involving me in decision-making | 8% |
| Being on time | 7% |
| Spending adequate time with me | 6% |

| Personal Responsibility N=226 | Percent Respondents |
| --- | --- |
| Exercise, diet and lead a healthy lifestyle | 47% |
| Shared decision making<sup>6</sup> | 35% |
| Follow medical recommendations given | 18% |

| Tests and Procedures N=226 | Percent Respondents |
| --- | --- |
| Shared decision making | 50% |
| I want all the tests that could be helpful to understand my condition better | 43% |
| I only want the absolute critical tests to be performed | 7% |

| Medications N=173* | Percent Respondents |
| --- | --- |
| Shared decision making<sup>7</sup> | 80% |
| I want the absolute minimum that I need to take for my condition | 9% |
| I want to take anything that can possibly help my condition | 9% |
| I want the freedom to try alternative medicine and herbal supplements | 2% |

| Healthcare Cost N=132** | Percent Respondents |
| --- | --- |
| I want to know what my health insurance covers | 57% |
| I want to know exactly what I am being charged for | 32% |
| I want to minimize my healthcare expenditure | 11% |

Combines respondents who picked the survey options “Kindness” and “Efforts to connect with me as a human being and not just as a patient”.
<sup>2</sup> Survey option was “Learn as much as I can about my condition and be actively involved in decision making”.
<sup>3</sup> Combines respondents who picked the survey options “I want to know exactly what I am taking and why” and “I want to understand the side effects of each medication thoroughly before accepting the prescription”.
\*Excluding 53 participants who were provided 4 choices instead of 5 choices for Question #4
\*\*Excluding 95 participants who were provided 4 choices instead of 3 choices for Question #5

Table 3 shows the top three most frequently selected qualities for each question.^4^ For question one assessing patient-physician relationship, humanistic qualities was the most frequent first (33%), second (24%) and third (30%) choice. For question number two assessing personal responsibility, healthy lifestyle (47%) was the most frequent top choice while learning about condition (38%) was the most frequent second choice. For tests and procedures, understanding the importance of diagnostic tests was the most frequent first (50%) and second (39%) choice. On question four assessing medications, understanding indication and side effects of medications was the most frequent first (80%), second (57%) and third (41%) choice. For question five on healthcare cost, knowing what insurance covers (58%) was the most frequent first choice while understanding charges (43%) was the most frequent second choice.

**Table 3.**
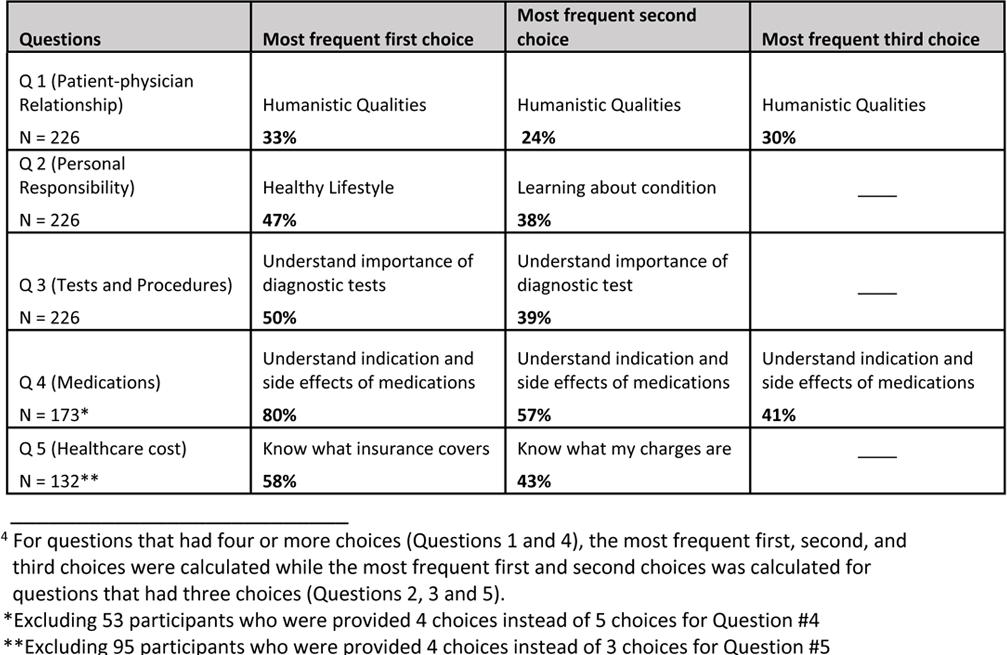
Top 1-3 qualities selected by respondents^4^

| Questions | Most frequent first choice | Most frequent second choice | Most frequent third choice |
| --- | --- | --- | --- |
| Q 1 (Patient-physician Relationship)<br>N = 226 | Humanistic Qualities<br>**33%** | Humanistic Qualities<br>**24%** | Humanistic Qualities<br>**30%** |
| Q 2 (Personal Responsibility)<br>N = 226 | Healthy Lifestyle<br>**47%** | Learning about condition<br>**38%** | _____ |
| Q 3 (Tests and Procedures)<br>N = 226 | Understand importance of diagnostic tests<br>**50%** | Understand importance of diagnostic test<br>**39%** | _____ |
| Q 4 (Medications)<br>N = 173* | Understand indication and side effects of medications<br>**80%** | Understand indication and side effects of medications<br>**57%** | Understand indication and side effects of medications<br>**41%** |
| Q 5 (Healthcare cost)<br>N = 132** | Know what insurance covers<br>**58%** | Know what my charges are<br>**43%** | _____ |

<sup>4</sup> For questions that had four or more choices (Questions 1 and 4), the most frequent first, second, and third choices were calculated while the most frequent first and second choices was calculated for questions that had three choices (Questions 2, 3 and 5).
\*Excluding 53 participants who were provided 4 choices instead of 5 choices for Question #4
\*\*Excluding 95 participants who were provided 4 choices instead of 3 choices for Question #5

Table 4 shows univariate analysis for demographic predictors of the most frequent top choice for each question (Q1: “humanistic qualities”; Q2: “healthy lifestyle”; Q3: “shared decision making”; Q4: “shared decision making” and Q5: “knowing insurance coverage”). When assessing patient-physician relationship, patients with college and above degrees and those with private insurance were less likely to rank humanistic qualities as their top choice compared to their references (0.55, CI 0.31-0.98 and 0.26, CI 0.11-0.64, respectively). For question two assessing patient personal responsibility, those 45 and older were less likely to rank healthy lifestyle as their number one choice when compared to those younger than 35 (0.20, CI 0.10-0.41 and 0.29, CI 0.11-0.77). Participants who identified their race as “Other”, those who had a college and above education and privately insured patients had higher odds of ranking healthy lifestyle as their number one choice compared to their references (2.55, CI 1.11-5.87; 4.25, CI 2.28-7.91 and 8.42, CI 2.42-29.33, respectively). When assessing preferences on tests and procedures, shared decision making was less likely to be ranked as a number one choice by those older than 65 (0.35, CI 0.13-0.99) but more likely to be ranked as a top choice by those with college and above education (2.01, CI 1.14-3.55) when compared to their respective references. With regards to medications, those who identified their race as “Other” had lower odds of choosing shared decision making as their top choice when compared to Blacks (0.24, CI 0.08-0.71). We saw no significant predicators for question five that assessed healthcare cost.

**Table 4.**
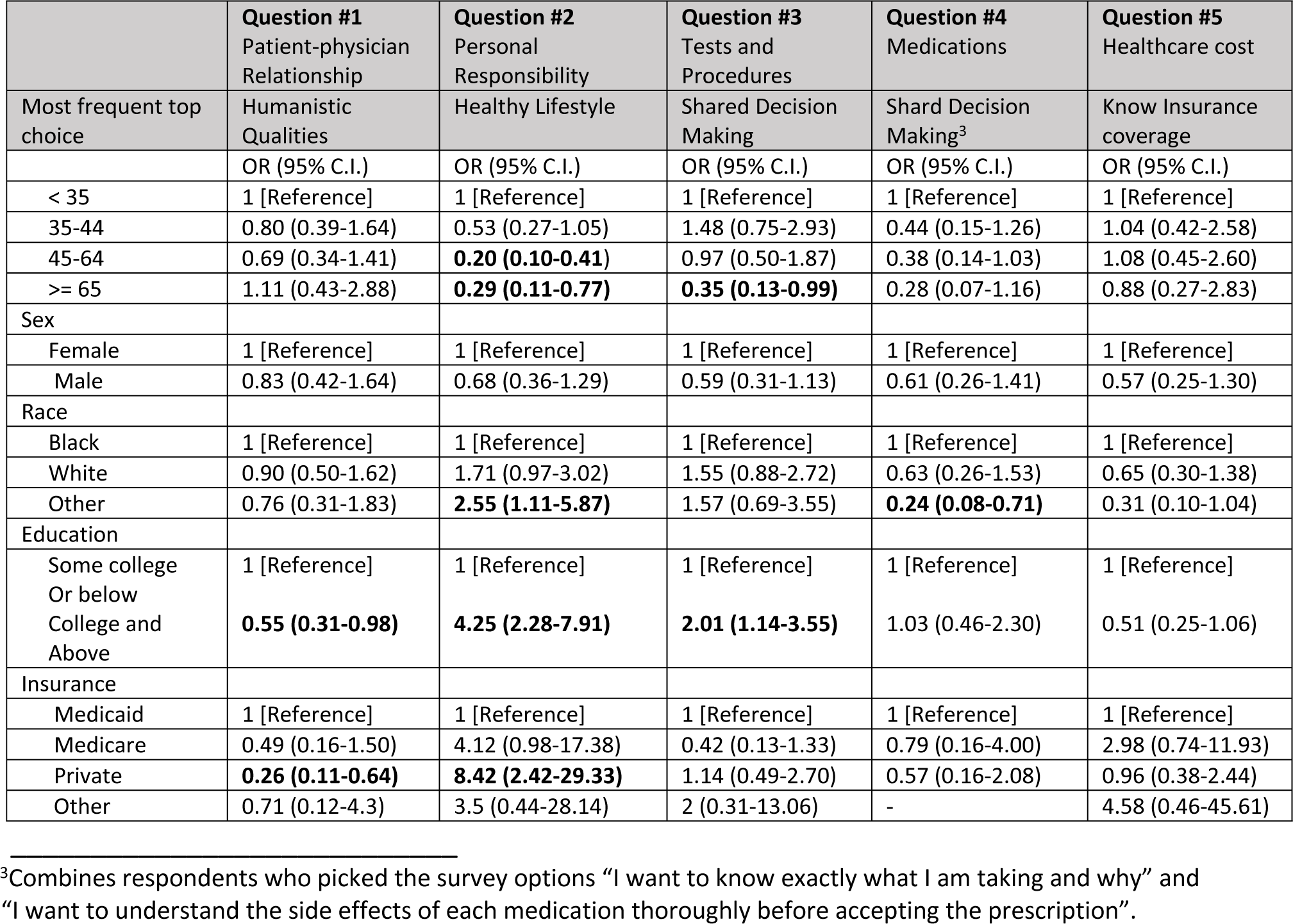
Univariate analysis for Predictors of Most Frequent Top Choice for Each Question

|  | Question #1<br>Patient-physician<br>Relationship | Question #2<br>Personal<br>Responsibility | Question #3<br>Tests and<br>Procedures | Question #4<br>Medications | Question #5<br>Healthcare cost |
| --- | --- | --- | --- | --- | --- |
| Most frequent top<br>choice | Humanistic<br>Qualities | Healthy Lifestyle | Shared Decision<br>Making | Shared Decision<br>Making<sup>3</sup> | Know Insurance<br>coverage |
|  | OR (95% C.I.) | OR (95% C.I.) | OR (95% C.I.) | OR (95% C.I.) | OR (95% C.I.) |
| < 35 | 1 [Reference] | 1 [Reference] | 1 [Reference] | 1 [Reference] | 1 [Reference] |
| 35-44 | 0.80 (0.39-1.64) | 0.53 (0.27-1.05) | 1.48 (0.75-2.93) | 0.44 (0.15-1.26) | 1.04 (0.42-2.58) |
| 45-64 | 0.69 (0.34-1.41) | **0.20 (0.10-0.41)** | 0.97 (0.50-1.87) | 0.38 (0.14-1.03) | 1.08 (0.45-2.60) |
| >= 65 | 1.11 (0.43-2.88) | **0.29 (0.11-0.77)** | **0.35 (0.13-0.99)** | 0.28 (0.07-1.16) | 0.88 (0.27-2.83) |
| Sex |  |  |  |  |  |
| Female | 1 [Reference] | 1 [Reference] | 1 [Reference] | 1 [Reference] | 1 [Reference] |
| Male | 0.83 (0.42-1.64) | 0.68 (0.36-1.29) | 0.59 (0.31-1.13) | 0.61 (0.26-1.41) | 0.57 (0.25-1.30) |
| Race |  |  |  |  |  |
| Black | 1 [Reference] | 1 [Reference] | 1 [Reference] | 1 [Reference] | 1 [Reference] |
| White | 0.90 (0.50-1.62) | 1.71 (0.97-3.02) | 1.55 (0.88-2.72) | 0.63 (0.26-1.53) | 0.65 (0.30-1.38) |
| Other | 0.76 (0.31-1.83) | **2.55 (1.11-5.87)** | 1.57 (0.69-3.55) | **0.24 (0.08-0.71)** | 0.31 (0.10-1.04) |
| Education |  |  |  |  |  |
| Some college<br>Or below<br>College and<br>Above | 1 [Reference]<br>**0.55 (0.31-0.98)** | 1 [Reference]<br>**4.25 (2.28-7.91)** | 1 [Reference]<br>**2.01 (1.14-3.55)** | 1 [Reference]<br>1.03 (0.46-2.30) | 1 [Reference]<br>0.51 (0.25-1.06) |
| Insurance |  |  |  |  |  |
| Medicaid | 1 [Reference] | 1 [Reference] | 1 [Reference] | 1 [Reference] | 1 [Reference] |
| Medicare | 0.49 (0.16-1.50) | 4.12 (0.98-17.38) | 0.42 (0.13-1.33) | 0.79 (0.16-4.00) | 2.98 (0.74-11.93) |
| Private | **0.26 (0.11-0.64)** | **8.42 (2.42-29.33)** | 1.14 (0.49-2.70) | 0.57 (0.16-2.08) | 0.96 (0.38-2.44) |
| Other | 0.71 (0.12-4.3) | 3.5 (0.44-28.14) | 2 (0.31-13.06) | - | 4.58 (0.46-45.61) |

<sup>3</sup>Combines respondents who picked the survey options “I want to know exactly what I am taking and why” and “I want to understand the side effects of each medication thoroughly before accepting the prescription”.

Table 5 shows multivariate analysis of demographic predictors of the most frequent top choice for each question (Q1: “humanistic qualities”; Q2: “healthy lifestyle”; Q3: “shared decision making”; Q4: “shared decision making” and Q5: “knowing insurance coverage”).The lower odds of choosing “humanistic qualities” associated with private insurance compared to having Medicaid persisted here (0.21, CI 0.07-0.65) but higher education dropped out when controlling for all other demographic characteristics. In the personal responsibility domain, higher odds associated with private insurance and higher education (5.73, CI 1.36-24.27 and 2.98, CI 1.34-6.59 respectively) as well as the lower odds of older age choosing healthy lifestyle (0.23, CI 0.11-0.51 and 0.32, CI 0.10-0.97) persisted compared to their reference groups respectively. When controlled for other factors, having “Other” race dropped out as a significant predictor whereas being on Medicare appeared to have significantly higher odds of association with choosing a healthy lifestyle compared to the Medicaid insured, although still with a very wide Confidence Interval (5.98, CI 1.24-28.93). For tests and procedures, having college and above education remained associated with higher odds of choosing SDM (2.30, CI 1.06-4.99) compared to lower educational attainment. In the Medication category having “Other” race persisted as having higher odds of choosing SDM compared to Blacks (0.16, CI 0.04-0.61). The healthcare cost category remained without significant association with any of the demographics we tested in both uni and multi-variate analyses.

**Table 5.**
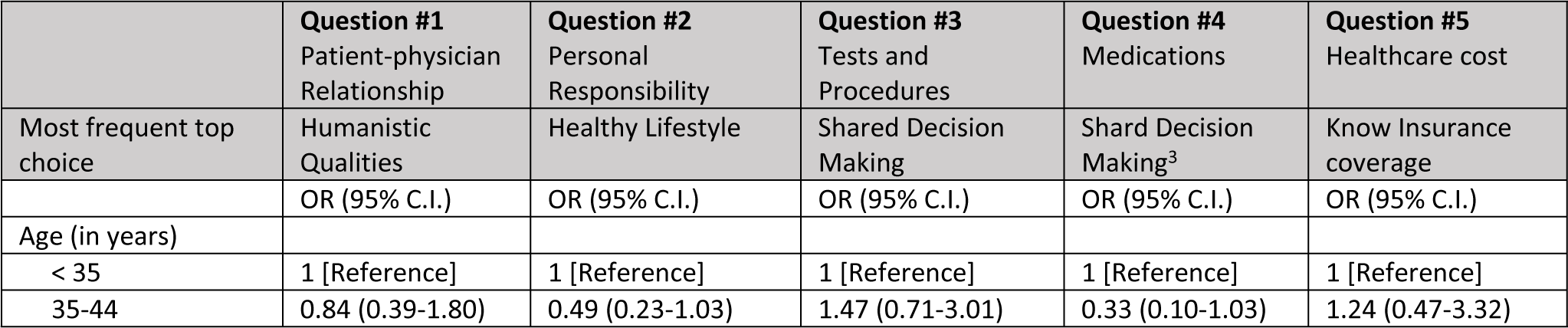
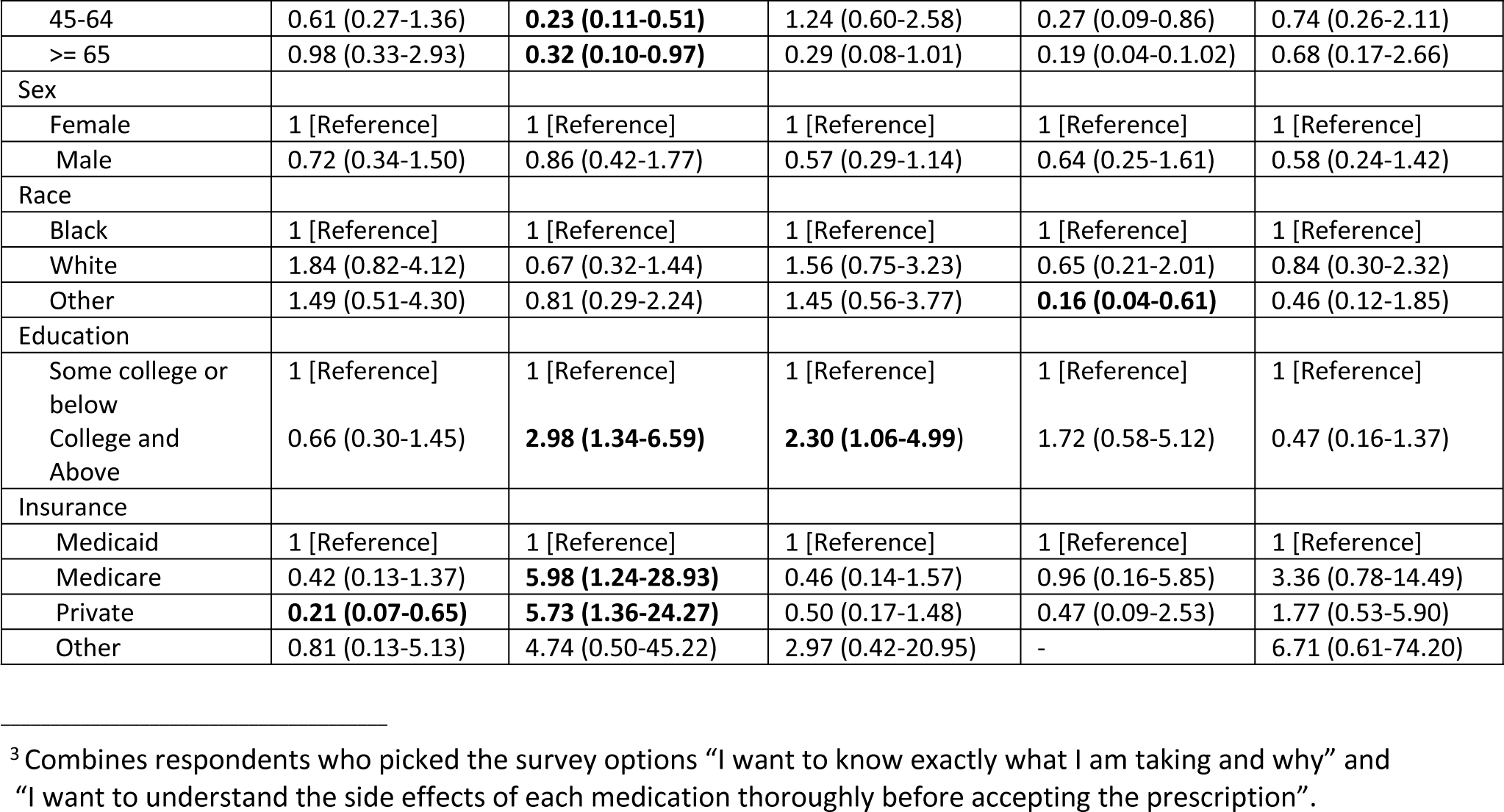
Multivariate Analysis for Predictors of Most Frequent Top Choice for Each Question

|  | **Question #1**<br>Patient-physician<br>Relationship | **Question #2**<br>Personal<br>Responsibility | **Question #3**<br>Tests and<br>Procedures | **Question #4**<br>Medications | **Question #5**<br>Healthcare cost |
| --- | --- | --- | --- | --- | --- |
| Most frequent top<br>choice | Humanistic<br>Qualities | Healthy Lifestyle | Shared Decision<br>Making | Shared Decision<br>Making<sup>3</sup> | Know Insurance<br>coverage |
|  | OR (95% C.I.) | OR (95% C.I.) | OR (95% C.I.) | OR (95% C.I.) | OR (95% C.I.) |
| Age (in years) |  |  |  |  |  |
| < 35 | 1 [Reference] | 1 [Reference] | 1 [Reference] | 1 [Reference] | 1 [Reference] |
| 35-44 | 0.84 (0.39-1.80) | 0.49 (0.23-1.03) | 1.47 (0.71-3.01) | 0.33 (0.10-1.03) | 1.24 (0.47-3.32) |
| 45-64 | 0.61 (0.27-1.36) | **0.23 (0.11-0.51)** | 1.24 (0.60-2.58) | 0.27 (0.09-0.86) | 0.74 (0.26-2.11) |
| >= 65 | 0.98 (0.33-2.93) | **0.32 (0.10-0.97)** | 0.29 (0.08-1.01) | 0.19 (0.04-0.1.02) | 0.68 (0.17-2.66) |
| Sex |  |  |  |  |  |
| Female | 1 [Reference] | 1 [Reference] | 1 [Reference] | 1 [Reference] | 1 [Reference] |
| Male | 0.72 (0.34-1.50) | 0.86 (0.42-1.77) | 0.57 (0.29-1.14) | 0.64 (0.25-1.61) | 0.58 (0.24-1.42) |
| Race |  |  |  |  |  |
| Black | 1 [Reference] | 1 [Reference] | 1 [Reference] | 1 [Reference] | 1 [Reference] |
| White | 1.84 (0.82-4.12) | 0.67 (0.32-1.44) | 1.56 (0.75-3.23) | 0.65 (0.21-2.01) | 0.84 (0.30-2.32) |
| Other | 1.49 (0.51-4.30) | 0.81 (0.29-2.24) | 1.45 (0.56-3.77) | **0.16 (0.04-0.61)** | 0.46 (0.12-1.85) |
| Education |  |  |  |  |  |
| Some college or below | 1 [Reference] | 1 [Reference] | 1 [Reference] | 1 [Reference] | 1 [Reference] |
| College and Above | 0.66 (0.30-1.45) | **2.98 (1.34-6.59)** | **2.30 (1.06-4.99)** | 1.72 (0.58-5.12) | 0.47 (0.16-1.37) |
| Insurance |  |  |  |  |  |
| Medicaid | 1 [Reference] | 1 [Reference] | 1 [Reference] | 1 [Reference] | 1 [Reference] |
| Medicare | 0.42 (0.13-1.37) | **5.98 (1.24-28.93)** | 0.46 (0.14-1.57) | 0.96 (0.16-5.85) | 3.36 (0.78-14.49) |
| Private | **0.21 (0.07-0.65)** | **5.73 (1.36-24.27)** | 0.50 (0.17-1.48) | 0.47 (0.09-2.53) | 1.77 (0.53-5.90) |
| Other | 0.81 (0.13-5.13) | 4.74 (0.50-45.22) | 2.97 (0.42-20.95) | - | 6.71 (0.61-74.20) |

<sup>3</sup> Combines respondents who picked the survey options “I want to know exactly what I am taking and why” and “I want to understand the side effects of each medication thoroughly before accepting the prescription”.

## Discussion

Our study showed that in the patient-physician domain, humanistic quality was the most frequently ranked top 1-3 choice. This is consistent with other research findings which document that patient-physician interaction is viewed by most patients to be a highly important aspect of quality care.[4,17,23] The higher value our patient population seems to place on their physicians’ humanistic over technical qualities such as the physician’s fund of knowledge could be explained by the fact that this survey was conducted in the ambulatory setting where a higher proportion of patients who may require significant emotional support are seen, an association that has been documented before.[11,19] Another possible reason why our patients showed a stronger preference for humanistic quality over technical quality is that patients who come to reputable healthcare settings may assume that they will be cared for by practitioners with superior technical abilities and hence tend to focus on their humanistic qualities instead.[37]

Although humanistic qualities appeared to be a highly valued choice for the domain of patient-physician relationship across the board, our uni and multi-variate analyses did show that the odds of choosing humanistic qualities was much lower for patients who had higher educational level (OR 0.55, CI 0.31-0.98) and or who were privately insured (OR 0.26, CI 0.11-0.64) as compared to lower educational level and Medicaid insured, a finding that has been noted before.[11,31] This may suggest that patients from lower socio-economic standing may have reasons to prioritize humanistic qualities in their care providers either because they don’t typically encounter this quality or because they may have increased needs for it due to their life circumstances.

In the personal responsibility domain, our findings of high correlation between prioritizing exercise, diet and leading a healthy lifestyle over other qualities with younger age, and higher educational attainment has been noted before.[38–41] This may be explained by the fact that, younger people are more agile, and a higher socio-economic standing (implied by higher educational attainment) may afford better access to healthy amenities as well as the fact that higher socio-economic standing may also confer the psychological space needed for people to prioritize healthy lifestyle over other concerns that may be at the top of their mind.[42–46] The higher odds of choosing healthy lifestyle seen in our multivariate analysis for those who have Medicare and Private insurance compared to the Medicaid insured (OR 5.98, CI 1.24-28.9 and OR 5.73, CI 1.36-24.27) is a significant finding and may once again be related to access to amenities in our patient population though this conclusion may carry less certainty for general interpretation due to the high confidence intervals.

Shared decision making (SDM) was the most frequently ranked top 1-3 choice for both “tests and procedures” and “medications” domains. The strong patient preference for SDM we found confirms the similar finding that has been noted before.[47,48] Clinicians will need to pay more attention to this aspect of care in the future as they will begin to see better informed patients come prepared to engage in decision making rather than to passively receive physician recommendations. The higher odds of choosing SDM by those with higher education in Q3 is also consistent with the evidence that better informed patients are likely to value and engage in SDM.[49,50] Explaining the higher odds we saw for choosing SDM for Q4 among those with “other” race would require a sub-subgroup analysis that was not performed here. In addition, loss of 53 surveys in this question may have reduced the power to detect other potentially significant associations in this category.

A question that asks patients to indicate their preference for knowing what their insurance covers and one that asks their preference to knowing what they are being charged for (two choices for Q5) is potentially confusing as one choice could be seen as a subset of the other. Despite that, it is clear that virtually all patients do care about the cost of care, especially the portion covered by insurance and/or themselves. Only a minority of those surveyed (11%) prioritized minimizing their healthcare expenditure which may indicate a related concept to the common health economics observation of moral hazard-where insured patients (virtually all our patients) may typically lack an incentive to prioritize healthcare cost minimization.

Our study has some weaknesses. The survey is liable to the inherent weakness of developing similar surveys discussed in the introduction. We attempted to mitigate some of that by designing it using similar survey concepts published previously,[33] and piloting the instrument before rolling out the project. Our survey population was mostly privately insured females which may limit the generalizability of some of our findings, but we have demonstrated statistically significant results from our logistic regressions that is worth replicating in a larger study to evaluate these findings. Incomplete and inaccurate completion of surveys that were excluded may have caused selection bias in our patient samples in addition to reducing our power in the analysis of results. Several patients who erroneously completed the surveys ranked multiple choices equally. Although this may be due to our survey design needing more clarity (as in for Q5) one of the inherent difficulty of accurately capturing patient priorities is their unwillingness to trade between quality attributes, a finding seen in studies with discrete choice experiments.[11] Given the move towards a patient centered model of care delivery, it will be important for the future to develop a validated instrument that captures what matters to patients in different settings.

Our study contributes to the growing body of evidence that patient centeredness and understanding patient priorities are essential for value-based care. Our findings are in line with other published studies that suggest that humanistic qualities,[4,17,23] healthy lifestyle,[38–41] and shared decision making[47,48] are important. In addition, our results extend what is known by showing that patients still prioritize these qualities even when offered equally attractive alternatives, and these priorities are associated with certain patient level factors.

In conclusion, the delivery of effective and quality medical care requires understanding of what most matters to patients. The task of deciphering the multiple factors that may affect patient priorities for what they value is a real challenge and may be criticized for having biases related to wording and context.[16] However, it is still a useful endeavor that can help clarify further what we may be able to achieve in our move towards a Value Based Care model that incorporates patients’ experience.

## Acknowledgments

We would like to thank all patient participants who were surveyed. We also thank Leah Jager, PhD for assistance with statistical analysis and Rachael Lebo for assistance with literature review.

